# Domain general frontoparietal regions show modality-dependent coding of auditory and visual rules

**DOI:** 10.1101/2024.03.04.583318

**Authors:** J. B. Jackson, A. N. Rich, D. Moerel, L. Teichmann, J. Duncan, A. Woolgar

## Abstract

A defining feature of human cognition is our ability to respond flexibly to what we see and hear, changing how we respond depending on our current goals. In fact, we can rapidly associate almost any input stimulus with any arbitrary behavioural response. This remarkable ability is thought to depend on a frontoparietal “multiple demand” circuit which is engaged by many types of cognitive demand and widely referred to as domain general. However, it is not clear how responses to multiple input modalities are structured within this system. Domain generality could be achieved by holding information in an abstract form that generalises over input modality, or in a modality-tagged form, which uses similar resources but produces unique codes to represent the information in each modality. We used a stimulus-response task, with conceptually identical rules in two sensory modalities (visual and auditory), to distinguish between these possibilities. Multivariate decoding of functional magnetic resonance imaging data showed that representations of visual and auditory rules recruited overlapping neural resources but were expressed in modality-tagged non-generalisable neural codes. Our data suggest that this frontoparietal system may draw on the same or similar resources to solve multiple tasks, but does not create modality-general representations of task rules, even when those rules are conceptually identical between domains.

## Introduction

Our sensory environment holds an abundance of information. This information is partly processed in specialised neural structures whose architecture supports a particular domain of processing, for example, primarily auditory input. However, flexible human cognition is generally thought to also require domain-general processing areas, capable of representing and integrating inputs from multiple modalities. Candidate domain general areas of the brain have long been identified (Duncan & Owen, 2000; Fedorenko, Duncan, & Kanwisher, 2013), but we do not know much about how information from multiple modalities is processed in these regions. For example, it is not clear to what extent cells in domain general regions exhibit preferences for sensory input from one modality over another, and to what extent they can be fully re-allocated to code different types of sensory information. Moreover, even if the neural resources are shared, it is not known to what extent information is represented in a similar or different way for each modality. In principle, the same information, arising from two different modalities, could be represented abstractly, with shared underlying neural codes, or independently, with non-generalisable patterns.

The frontal-parietal multiple-demand (MD) network is a functionally integrated neural circuit that is recruited by many types of cognitive demand (Duncan & Owen, 2000; Duncan, 2010; Fedorenko et al., 2013; Assem, Shashidhara, Glasser, & Duncan, 2022). It is thought to play a key role in cognitive control by integrating the relevant information from multiple more-specialised systems, as needed for the current cognitive operation (Cole & Schneider, 2007; Cole et al., 2013; Cocuzza, Ito, Schultz, Bassett, & Cole, 2020; Duncan, Assem, & Shashidhara, 2020). The regions of this system show high correlations of their functional timeseries both with and in the absence of a cognitive task (Power et al., 2011; Ji et al., 2019; Cocuzza et al., 2020), and co-activate during different demanding tasks including those associated with working memory, selective attention, and problem solving (Fedorenko et al., 2013; Assem, Glasser, Van Essen, & Duncan, 2020; Shashidhara, Spronkers, & Erez, 2020). Single-unit work in non-human primates has shown that information from different modalities is represented within this network, with prefrontal and parietal neurons encoding information about both auditory (Azuma & Suzuki, 1984; Romanski, 2007) and visual stimuli (Rao, Rainer, & Miller, 1997; Freedman, Riesenhuber, Poggio, & Miller, 2001; Freedman & Assad, 2006). Similarly in human brain studies, the MD network codes a variety of task features including task relevant information from tactile (Woolgar & Zopf, 2017), visual, auditory, rule, and motor domains (Woolgar, Jackson, & Duncan, 2016; Pischedda, Görgen, Haynes, & Reverberi, 2017; Vaidya, Jones, Castillo, & Badre, 2021; Schultz, Ito, & Cole, 2022), consistent with the idea of a domain general network.

Within the domain of vision, we know that MD responses are flexible. For example, the representation of visual and rule information in these regions adjusts according to task demands (Woolgar, Hampshire, Thompson, & Duncan, 2011; Woolgar, Afshar, Williams, & Rich, 2015). Visual studies have also shown that these regions exhibit preferential coding for attended over unattended objects (e.g., Woolgar, Williams, & Rich, 2015) and relevant over irrelevant dimensions of visual objects (Jackson, Rich, Williams, & Woolgar, 2017; Jackson & Woolgar, 2018). The neural patterns in these MD regions appear to be important for behaviour (Woolgar, Dermody, Afshar, Williams, & Rich, 2019; Robinson, Rich, & Woolgar, 2022) and causal for flexible coding elsewhere in the network (Jackson, Feredoes, Rich, Lindner, & Woolgar, 2021). We have previously shown that codes for different visual stimuli are held by overlapping MD voxels (Jackson & Woolgar, 2018). In addition, recent work using high-precision functional magnetic resonance imaging (fMRI) has reported almost identical activation maps for hard compared to easy versions of auditory and visual tasks, even at the single-subject level (Assem et al., 2022). These studies suggest that resources in the MD network can be flexibly allocated to represent relevant task information.

Based on this prior literature, the MD network appears to be domain general, flexibly representing relevant rules and stimuli regardless of the source modality. However, there are different possibilities for how information from distinct modalities is structured within these regions. This is important because there are several possible conceptualisations of domain generality. First, at the most basic level, a network could be considered domain general merely because it responds to multiple inputs. This aligns with the original observation of the MD network (Duncan & Owen, 2000; Fedorenko et al., 2013), which is defined as a network that responds during different cognitively challenging tasks. Such a result could arise if activity in these regions reflected task-general processes such as effort or error monitoring, common to many tasks. Second, a stronger requirement for domain generality might be that the network not only responds to, but also represents, multiple types of information. For example, it could show specific patterns of activity from which it is possible to decode the details of, for example, both visual and auditory stimuli (Woolgar et al., 2016). This could arise because the network comprises dedicated resources responding to each modality-specific task that are co-located in the same network or regions (Buschman, 2021), giving rise to generality at the region or network, but not the single cell, level. Third, domain generality could be defined as when a network re-uses the *same resources* to process these different types of information in different situations. This could arise if general-purpose resources are flexibly allocated to process information from each of the two modality-specific tasks (for example, as in Manohar, Zokaei, Fallon, Vogels, & Husain, 2019; Bocincova, Buschman, Stokes, & Manohar, 2022). This is challenging to assess with fMRI (as even our highest resolution protocols capture the activity of tens of thousands of neurons) but we can at least ask whether patterns from multiple modalities load onto the same or different voxels (Jackson & Woolgar, 2018). Finally, a network showing flexible re-use of general-purpose resources may express this domain generality in different ways. It could hold information in an abstract form that generalises over input modality, or in a modality-tagged form, which re-uses the same resources but utilises unique codes to represent the information in each modality. In the former case, the same resources would be used in the same way between modalities, and in the latter, the same resources may be used, but in different ways. Information about the input modality could be preserved in either scheme, in the former by inclusion of a separate modality-signalling code (an additional dimension in the representational scheme (Badre, Bhandari, Keglovits, & Kikumoto, 2021)), and in the latter by changing the content code sufficiently between modalities so that codes do not generalise.

Here, we distinguished between these possibilities by considering how information from two different sensory modalities is represented in the MD network. Participants applied identical stimulus-response mapping rules to visual or auditory stimuli. Using multivariate pattern analysis (MVPA) to characterise the MD representation of these rules, we asked 1) whether the MD network codes for rule information for both auditory and visual tasks; 2) whether the same voxels contribute to both sets of codes; and 3) whether the codes underlying these rule representations are modality-specific, or abstract and modality-independent. This allowed us to assess whether visual and auditory task rules are represented in this network by independent codes, or whether MD representation of task rules is abstracted away from the input modality, with codes that generalise between auditory and visual tasks. Aligning with the concept of the MD system as a flexible resource, but against the intuition that conceptually identical rules should be represented in an abstract, generalisable form, our results show strongly decodable rules in the two modalities that draw on overlapping resources but are represented in modality-tagged non-generalisable codes.

## Methods

### Participants

We recruited 49 volunteers for the experiment. Of these, 17 were excluded for not passing the training or for not completing all the sessions (2 behavioural training sessions plus 1 fMRI session). The final cohort therefore consisted of 32 participants (mean age 24 years, SD = 4.1 years; 2 left-handed; 9 men, 24 women). Participants were required to pass MRI safety screening, have normal or corrected-to-normal vision and no history of neurological or psychiatric disorder. Participants were recruited through word of mouth and from the Macquarie University SONA participation pool at the Department of Cognitive Science, Macquarie University. They gave written informed consent to participate and were compensated for their time. The experiment was approved by the University of Macquarie Research Ethics Committee (reference: 5201300541).

### Task design

Participants completed a task with visual or auditory stimuli (based on Jiang & Kanwisher, 2003), with trials from the two modalities randomly interleaved (Figure 1a). Before the start of each trial a tone was played, and simultaneously a white cross was displayed on the screen (500ms) followed by a black cross (500ms) to indicate the upcoming trial. At the start of each trial, the trial type (auditory/visual), and current stimulus-response mapping rule, was indicated by a word (auditory trials) or a symbol (visual trials) for 540ms. In the auditory trials, participants then heard four consecutive tones (200ms each, 200ms between tones) and had to identify which tone (1-4) had the highest pitch. A black fixation cross was displayed throughout the auditory trials. In the visual trials, participants observed 4 consecutive vertical lines (200ms each, 200ms between displays) and had to identify which was shortest (1-4). Participants responded by pressing one of four response buttons using the index and middle fingers on both hands. There were two different stimulus-response mapping rules (Figure 1b). Each rule had two cues per modality (e.g., for auditory trials for a specific participant, two words “dogs/post” for rule 1, and two words “milk/wish” for rule 2; cue-rule association counterbalanced over participants, see Figure 1c). This was to avoid the classifiers using activity related to the specifics of the cue to distinguish the rule conditions. There was an equal number of trial types in each run (modality, rule, cue, and target position; 32 trials, in total 6 runs and 192 trials).

**Figure 1.**
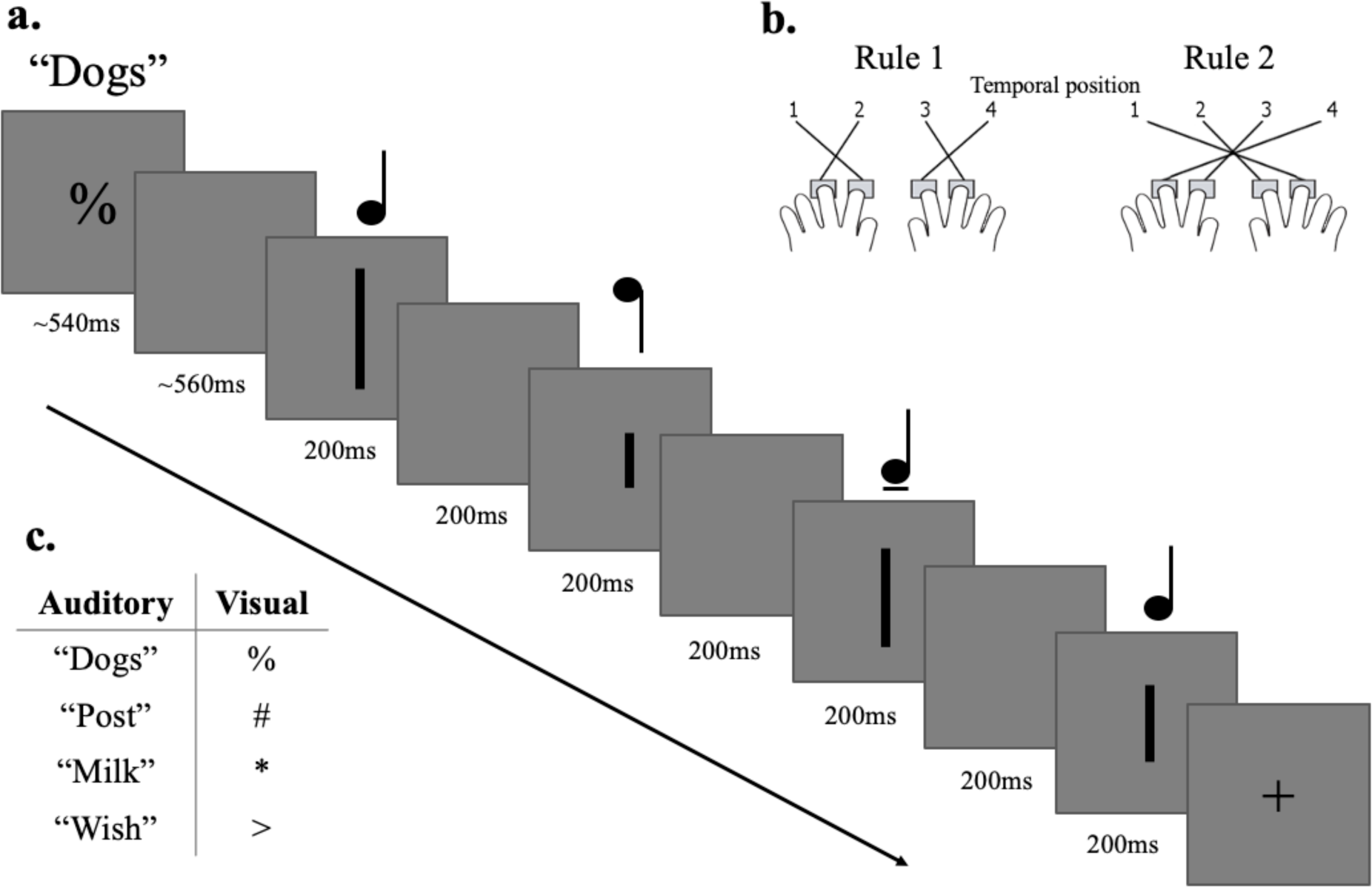
Task design. Panel a) depicts a trial from the main task. Participants were presented with a rule cue for 540ms. This was either a word (auditory task) or a symbol (visual task). In the auditory trials participants were asked which tone had the highest pitch out of four consecutive tones. A black fixation cross was presented on the screen through the duration of the auditory trial (not depicted here). In the visual trials, participants were asked which line was the shortest out of four consecutive vertical lines. Stimulus presentation was 200ms with a 200ms gap in between each presentation. Panel b) shows the two stimulus-response mappings (rules). For example, under rule 1, to indicate that the highest tone or shortest line was in the second temporal position (as shown), the participant should press the first (far left) button. Under rule 2, they should instead press the third button. Each rule had two cues per modality. The spoken words used for the rule cues are from Petit et al., (2020). Panel c) shows the cues for the auditory and visual modalities, which we assigned to the two rules in a manner that was counterbalanced over participants.

### Procedure

#### Overview

Participants completed three sessions in total. In the first two sessions (training, 1 hour each) participants completed titration and training for the main task. During these sessions, task difficulty was equated for the auditory and visual stimuli using a staircase procedure, and participants practiced the different stimulus-response mappings (rules). In the final session participants completed the main task in the scanner. Below we outline the procedure in more detail.

#### Session 1 and 2 (training)

The training in this first session had five separate stages. In the first stage participants practised identifying the temporal position of the highest tone or shortest line, without the stimulus-response mappings or associated rule cues, separately for the two task modalities. Participants were presented with a block of trials for each modality and responded by saying number 1-4, depending on which tone had the highest pitch (auditory) or which line was the shortest (visual). The experimenter would then enter the number reported. There were two blocks in total (20 trials). Participants did not respond using the keyboard to avoid them explicitly practicing a stimulus-response mapping that would conflict with the stimulus-response mapping used in the scanning session. Participants received feedback on every trial (fixation cross changed to green or red, 500ms) and there was no response time out. In all stages of the experiment the starting modality was counterbalanced by participant number.

In the second stage of training, participants repeated the first stage but this time with a staircase to equate task difficulty for the auditory and visual stimuli. We ran 2 (4-down, 1-up) staircases per modality (4 staircases total). The 4-down, 1-up staircase was chosen to ensure high task accuracy. The 4 staircases were blocked and each staircase ended after a maximum of 10 reversals. The threshold was calculated by averaging the threshold at the final reversals for each staircase (discarding the first 4 reversals if there was an even number of reversals, or discarding the first 3 if there was an odd number of reversals). The 2 threshold estimates per modality were then averaged. Participants received feedback on every trial (fixation cross changed to green or red, 500ms) and there was no response time out.

In the third stage of training, participants were shown the stimulus-response mappings and were required to enter responses themselves using the keyboard. Participants used the A, S, K, and L buttons on the keyboard to respond using the same fingers as in the fMRI task. Each stimulus-response mapping was introduced sequentially (e.g., rule 1 visual stimuli, rule 1 auditory stimuli, rule 2 visual stimuli, rule 2 auditory stimuli), and the participants learnt to associate the rule cues with each rule. The starting rule was selected pseudorandomly. Participants completed 4 blocks (12 trials per block) and after the first 2 trials of each block they were shown a reminder of the rule. Participants received feedback on every trial (fixation cross changed to green or red, 500ms) and there was a response time out after 6s.

In the fourth stage of training, participants practiced the task with the two rule cues combined and had to use the cues to differentiate which rule to apply. In this stage participants completed 4 blocks (40 trials in total). The rule-cue mapping was kept the same for each participant throughout all sessions of the experiment. In the final fifth stage, the modalities were also interleaved making the task very similar to that in the scanning session. Participants still received feedback in this fifth stage, and they completed a minimum of 80 trials.

At the start of the second training session participants were asked again to sign a behavioural consent form. They then practised the (stage 5) task until the time for that session had ended.

#### Session 3 (MRI)

In session 3, participants filled out an MRI consent and screening form. To remind them of the task, participants practised the task with feedback outside the scanner room (minimum of 80 trials). They then received instructions on the button box, and we acquired a structural scan. To get familiar with the scanner setup and button box participants practised the task in the scanner for 8 trials in total (with feedback). Following this they completed 6 runs of the main task without feedback (refer to section: Task design).

### Data acquisition

We collected the data with a Siemens (Erlangen, Germany) 3-T Verio scanner at Macquarie Medical Imaging, Macquarie University Hospital, Sydney, Australia. We acquired a T1-weighted structural image for each participant (1mm isotropic voxels, repetition time (TR) 2000ms, echo time (TE) 2.36ms). To avoid acoustic noise from the scanner during presentation of the auditory (and visual) stimuli, we used an interleaved steady state (ISSS) imaging sequence (Peelle, Eason, Schmitter, Schwarzbauer, & Davis, 2010; Peelle, 2014), with a delay of 1 TR (RF and pulsed steady state silent mode, with 60us ramp time) after every 4 volumes. Volumes were acquired using interleaved T2*-weighted EPI acquisition with the following parameters: TR 3000ms; TE 33ms; 32 slices of 4.5 mm slice thickness with no interslice gap; in-plane resolution 3.0 × 3.0mm; field of view 250mm.

### Preprocessing

We preprocessed the MRI data using SPM 12 (Wellcome Department of Imaging Neuroscience, www.fil.ion.ucl.ac.uk/spm) in MatLab 2018a. We converted functional MRI data from DICOM to NIFTI format and spatially realigned to the first functional scan. We did not perform slice-timing correction due to the non-continuous acquisition (ISSS sequence) (as in Perrachione & Ghosh, 2013). We co-registered structural images to the mean EPI. We smoothed the EPIs in-plane (in the x and y, but not in the z direction) with a 4 mm FWHM Gaussian kernel. We also high pass filtered (128s) the data. Finally, we normalised the structural scans to the T1 template of SPM12 (Wellcome Department of Imaging Neuroscience, London, UK; www.fil.ion.ucl.ac.uk), using SPM12’s segment and normalise routine. This was to derive the individual participant normalisation parameters needed for transformation of ROIs into native space and to normalise the searchlight classification maps derived in native space.

### ROIs

We took 13 frontal and parietal MD ROIs from the parcellated map provided by Fedorenko et al (2013; available online at imaging.mrc-cbu.cam.ac.uk/imaging/MDsystem). This map consists of regions that show increased activation with task demands across a range of tasks. This definition of the MD network shares a high degree of overlap with the previous definition (Duncan & Owen, 2000) derived from meta-analytic data, that we used in previous work (Woolgar, Afshar, et al., 2015; Woolgar, Williams, et al., 2015; Jackson et al., 2017; Jackson & Woolgar, 2018; Jackson et al., 2021). MD ROIs comprised left and right anterior inferior frontal sulcus (aIFS; centre of mass (COM) = ±35 47 19, volume = 5.0 cm^3^), left and right posterior inferior frontal sulcus (pIFS; COM ±40 32 27, 5.7 cm^3^), left and right premotor cortex (PM; COM ±28 −2 56, 9.0 cm^3^), left and right inferior frontal junction (IFJ; COM ±44 4 32, 10.1 cm^3^), left and right anterior insula/frontal operculum (AI/FO; COM ±34 19 2, 7.9 cm^3^), left and right intraparietal sulcus (IPS; COM ±29 −56 46, 34.0 cm^3^), and bilateral pre-supplementary motor area/anterior cingulate cortex (pre-SMA/ACC; COM 0 15 46, 18.6 cm^3^). For the main analyses we combined these ROIs into one MD network ROI using FSL v5.08 functions (Jenkinson, Beckmann, Behrens, Woolrich, & Smith, 2012). For decoding results from individual MD ROIs see supplementary materials.

We defined early visual cortex (BA17: COM −1 −79 6, 31 cm^3^) and primary motor cortex (BA4: COM ±27.8 −23 60, 51.6 cm^3^) from the Brodmann template provided with MRICroN (Rorden, 2007). We also defined primary auditory cortex (A1) as a spherical ROI (radius 10 mm) placed at the intersection of BA 41 and 42 within Heschl’s gyrus (COM ±50 −22 10, 31.9 cm^3^). The decoding results for the visual and auditory ROIs can be found in the supplementary materials.

All ROIs were deformed into native space by applying the inverse of the normalisation parameters for each participant.

### First-Level Model

To obtain activation patterns for MVPA, we estimated two separate GLMs for each participant (SPM12). For the main model, we estimated the activity associated with the two visual, and two auditory rules, using correct trials only (4 regressors). To account for trial by trial variation in reaction time (Todd, Nystrom, & Cohen, 2013), trials were modelled as events lasting from the offset of the fourth stimulus until response convolved with the hemodynamic response.

We then ran a second model designed to capture button press responses. We used this as a sanity check for whether we could successfully decode the given response (inner vs outer finger position) in each task modality separately (within-modality decoding), and whether this information could be cross-generalised across modality (between-modality decoding) in motor regions. For this we estimated the activity associated with the inner and outer finger responses across both hands, for the visual and auditory tasks separately (4 regressors total, modelled as events lasting from fourth stimulus offset until response convolved with the hemodynamic response).

For both GLMs we included dummy scans and movement parameters (translation and rotation) as covariates of no interest (totalling 7 regressors). We maintained the correct relationship between events and fMRI volumes by adding dummy volumes into the time series to occupy the silent time periods where no data were collected. As a reminder, the purpose of the silent time periods was to avoid acoustic noise from the scanner during stimulus presentation (see section Data acquisition). We then removed their influence on parameter estimation by perfectly modelling them with a single regressor (0 for actual volumes, 1 for dummy volumes) that was not convolved with the haemodynamic response function (as described in Peelle, 2014). As we did not perform slice-timing correction (ISSS sequence), we also estimated temporal derivatives to account for slice-time differences. Temporal derivatives were estimated for the main task (the visual and auditory rules in the first GLM, and the inner and outer finger responses in the second GLM) and for the dummy regressors. We combined the two parameter estimates (e.g. visual rule 1, and its temporal derivative) by the root mean squared (Calhoun, Stevens, Pearlson, & Kiehl, 2004) and took these estimates forward for our decoding analysis.

### Analysis

We used MVPA to examine the representation of the rules applied to the visual and auditory stimuli. Of central interest was 1) whether the MD regions coded both visual and auditory-based rule information; 2) whether the same voxels were re-used for both sets of codes; and 3) whether rule representations generalised across modality, suggesting modality-independent rule representations.

We implemented MVPA using the Decoding Toolbox (Hebart, Görgen, & Haynes, 2015). To address our first question of whether the MD regions coded both visual and auditory rule information, we trained the classifier to discriminate the two stimulus-response mappings (rules) for each modality separately (within-modality decoding). We did this in the combined MD ROI, in each MD ROI separately, and in the visual and auditory ROIs.

For our second question, we asked to what extent the same voxels contributed maximally to the classification of rule in each modality (auditory and visual). For this we extracted the transformed classifier weights for each classification scheme (visual rule/auditory rule). Raw classifier weights are not a simple reflection of the signal at each voxel, but an index of signal strength that can be recovered by multiplying the raw classifier weights by the data covariance (transformed classifier weights, Haufe et al., 2014). We identified the voxels with the highest (top 10%) of these signal-reflecting transformed classifier weights for visual rule coding and the top 10% of voxels with the highest transformed weights for auditory rule coding, and asked how many of these were the same voxels (expressed as a proportion, as in Jackson & Woolgar, 2018). We then used a two-step permutation test (Stelzer, Chen, & Turner, 2013) to test whether the proportion of voxels that contributed to both classification schemes in our data exceeded the proportion expected by chance. For each person, we trained a classifier on the permuted condition labels for each classification scheme (visual rule 1/visual rule 2, and auditory rule 1/auditory rule 2) which resulted in 32 unique combinations. We then built a voxel re-use null distribution for each participant by randomly selecting one of the 32 unique combinations from each classification scheme and calculating the proportion of overlap in the top 10% of voxels with the highest transformed weights. We ran 10,000 permutations of this. We then built a group level null distribution by sampling (with replacement) 1000 samples per participant from these permutations and then finally averaging the 1000 samples across participants. From the group null distribution, we then calculated the probability of observing the actual voxel re-use value by means of the Monte-Carlo approach (*p* = *k*+1/(*n*+1)) where *k* is the number of permutations in the null with equal or higher accuracy to the actual voxel re-use value and *n* is the number of all permutations. The estimate we used is slightly more conservative than the unbiased estimate (*p = k / n)* and avoids returning a *p* value of 0 (Phipson & Smyth, 2010; Hemerik & Goeman, 2018). Separate to this analysis we also averaged the transformed weights in each region of the MD network to depict the relative weighting, or contribution, of individual MD ROIs to the two classification schemes.

To address the third question of whether auditory and visual rule information cross-generalised, we trained a classifier to distinguish the visual rules (visual rule 1 vs visual rule 2) and tested performance on discriminating the auditory rules (auditory rule 1 vs auditory rule 2), and *vice versa*, training on the auditory rules and testing on the visual rules (between-modality decoding). Next, as a positive control and to assess whether we could distinguish between the codes representing the two modalities, we trained a classifier to discriminate between the auditory (rule 1 and rule 2 data) and visual task (rule 1 and rule 2 data). We performed these analyses in the combined MD network ROI, in each MD ROI separately, and in the visual and auditory ROIs. To check whether there were additional regions that showed cross-generalisation, we also performed an exploratory analysis in which we carried out classification across the whole brain using a roaming searchlight (Kriegeskorte, Goebel, & Bandettini, 2006). For each participant, data were extracted from a spherical ROI (radius 5mm) that was centred on each voxel in the brain. A classifier was trained and tested using data from each sphere, and the classification accuracies were assigned to the central voxel. This yielded whole-brain accuracy maps for each individual. Accuracy maps were normalised and smoothed (8mm FWHM Gaussian kernel) for group-level analysis (one-sample *t*-test at each voxel). The results were thresholded at *p* < 0.005 with an extent threshold of 20 voxels.

For the final classification schemes and as a sanity check, we assessed whether we could decode motor responses in the visual and auditory-based tasks separately (within-modality decoding), and whether these representations cross-generalised from the auditory to the visual task and *vice versa* (between-modality decoding). For this we trained the classifier to discriminate inner vs outer finger responses, training and testing within modality (visual/auditory), followed by training the classifier on inner vs outer responses in the visual task and testing the classifier on inner vs outer responses in the auditory task (and *vice versa*). We looked at information pertaining to these motor responses in the primary motor cortex (BA4).

For all classification analyses we used a linear support vector machine and a leave-one-out six-fold splitter. For the within-modality decoding analysis the classifier was trained using the data from five of six runs (e.g., for the visual modality: visual rule 1 vs rule 2) and subsequently tested on its accuracy at classifying the unseen data from the remaining run (visual rule 1 vs rule 2), iterating over all possible combinations of training and testing runs. For the between-modality decoding analysis the classifier was trained using the data from five of six runs from one modality (e.g., visual rule 1 vs rule 2, runs 1:5) and tested on its accuracy at classifying the left out run from the other modality (e.g., auditory rule 1 vs rule 2, run 6), iterating over all possibilities (and vice versa; train on auditory rules, test on visual rules). For our positive control to test if we could distinguish auditory from visual information, we again trained the classifier on five runs (visual rule 1 and 2 vs auditory rule 1 and 2) and tested on a final sixth run (visual rule 1 and 2 vs auditory rule 1 and 2) with all possible iterations. The accuracies were averaged to give a mean accuracy score. This was repeated for each condition, participant, and ROI separately.

We used Bayesian statistics (Kass & Raftery, 1995; Rouder, Speckman, Sun, Morey, & Iverson, 2009; Dienes, 2011; Morey, Romeijn, & Rouder, 2016) to determine the evidence for above-chance decoding (alternative hypothesis) and chance decoding (null hypothesis) for each classification scheme and ROI using the Bayes Factor (BF) R package (Morey et al., 2016). We used a half-Cauchy prior for the alternative hypothesis to capture directional (above chance) effects. The prior was centred around chance (δ = 0, i.e., 50% decoding accuracy), and had the default width of 0.707 (Jeffreys, 1998; Wetzels et al., 2011). We excluded the interval ranging δ = 0-0.5 from the prior to determine very small effect sizes as irrelevant (Teichmann, Moerel, Baker, & Grootswagers, 2021). We interpreted BFs below 1/3 or above 3 as evidence for the null or the alternative hypothesis respectively (Wetzels et al., 2011).

## Results

### Behavioural data

Reaction time and accuracy data from the scanning session are depicted in Figure 2. Participants performed with a high degree of accuracy (mean percent correct for the visual rules = 93.6%, *SD* = 4.9, mean percent correct for the auditory rules = 94.7%, *SD* = 5.3). There was evidence for no difference in accuracy scores between the two modalities at the group level (Bayesian paired t-test BF_10_ = 0.3). There was strong evidence that participants took longer to respond on the auditory task than the visual task (mean auditory RT = 1699.3ms, mean visual RT = 1165.6ms, Bayesian paired t-test BF_10_ = 391002.3). This was also the case for 28/32 participants at the single subject level.

**Figure 2.**
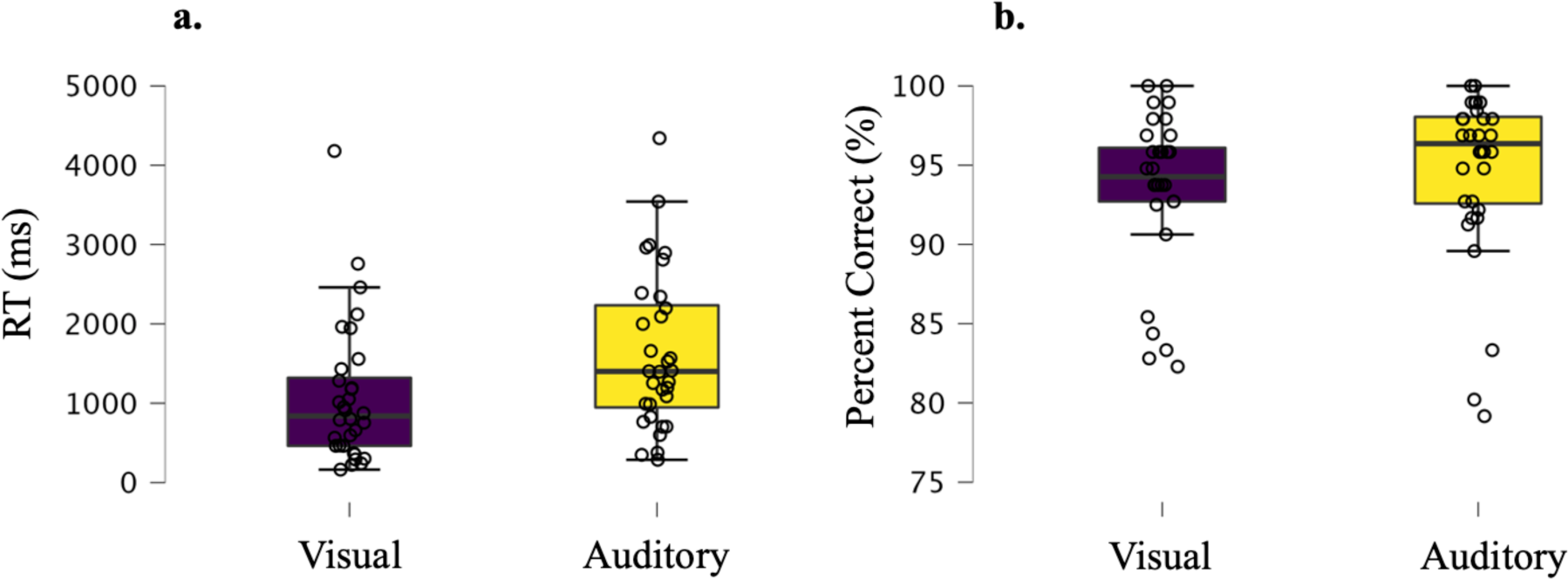
RT (correct trials only) and accuracy (percent correct, %) data. Boxplots (plotted in JASP (Team, 2024)) with individual data points (open circles) in Panel a) show RT (ms) data and Panel b) show accuracy data. Boxplot inner line depicts median, and the box spans the range from the 25^th^ to the 75^th^ percentile. The whiskers extend to the 25th percentile - (1.5* interquartile range) for the lower boundary, and 75th percentile + (1.5* interquartile range) for the upper boundary. If the value for either is 0 then the whiskers extend to the data extremes. Participants performed with a high degree of accuracy overall due to the extensive training and performance criteria for progressing to the fMRI session.

### Coding of visual and auditory-based rule information

To address our first question of whether the MD network codes rule information in both visual and auditory tasks, we used MVPA to differentiate the patterns pertaining to the visual (visual rule 1 vs visual rule 2) and the auditory rules (auditory rule 1 vs auditory rule 2) separately. The resulting classification accuracies and associated BFs are shown in Figure 3a. Rule information was encoded in the MD network (combined ROI) when participants completed the task in the visual (mean classification accuracy 58.4%, BF_10_ = 233.8) and the auditory domains (mean classification accuracy 57.4%, BF_10_ = 3.1). For individual MD ROI results refer to supplementary figure 1.

**Figure 3.**
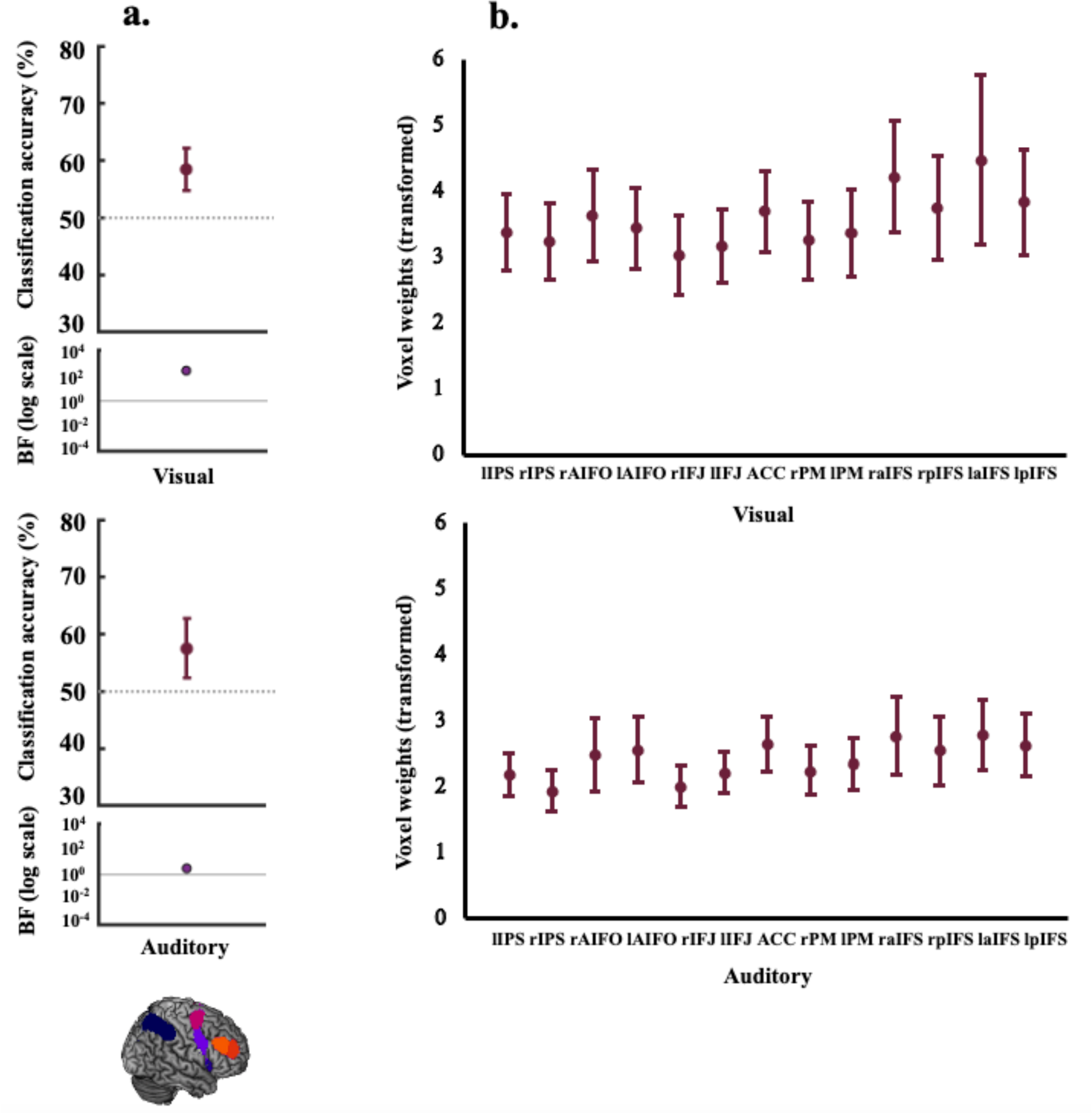
Decoding results and transformed weights for combined MD network ROI. Panel a) shows coding (classification accuracy, %) of visual rules (upper) and auditory rules (lower) (within-modality decoding). The lower parts of the plots show the associated BFs on a logarithmic scale, generated using custom code (from Teichmann et al., 2021). BF_10_ < 1/3 are marked in blue and BF_10_ > 3 are marked in purple-coloured circles. There was evidence to indicate that the MD network encoded rule information in both the auditory and visual tasks. Panel b) depicts a projection of the transformed weights (absolute) for each classification scheme (visual rules, auditory rules) averaged in the individual MD ROIs. These plots indicate that classification of the auditory and visual rules was not driven by any region of this network in particular. For plotting purposes, we bootstrapped 95% confidence intervals across participants using 10,000 bootstrap samples.

Next, we plotted the average of the transformed absolute classifier weights for each classification scheme (visual/auditory; Figure 3b) across the individual MD ROIs to assess their relative contributions. The weights were evenly distributed over ROIs, indicating that classification performance was not driven by any subset of ROIs.

### Voxel re-use across auditory and visual tasks

Thus far the results show that the MD network holds information pertaining to both the auditory and visual rules. However, it is possible that information about these two modalities is still carried by independent populations of neurons within this network. In this model, individual MD voxels might have structured preferences for rule coding in the two modalities. We examined the overlap (re-use) in the 10% of voxels that contributed the most signal to each classification scheme and ran permutation tests allowing us to compare the observed proportion of voxel re-use to that expected by chance. The MD network displayed a higher proportion of voxels used in both classification schemes (33.4%, *p* < .0001) than what would be expected by chance (26.9%, group null average), suggesting that the same voxels were used to encode information about both task modalities.

### Cross-generalisation of visual and auditory information

Our next question was whether the representation of rules cross-generalised between modalities, that is, whether the MD network codes for rule information in the same manner regardless of the input modality. Alternatively, it may be that while overlapping voxels in this network encode information about both modalities, the codes are not shared and are independent. Consistent with this latter possibility, the data presented in Figure 4a show evidence for the null that there was no cross-generalisation of auditory and visual rule information in this network. The mean classification accuracy for training on the visual rules and testing on the auditory rules was 48.4%, (BF_10_ = 0.16), and for training on auditory and testing on visual, it was 51.1%, (BF_10_ = 0.11). We further tested the complementary question of whether this network holds modality-distinguishing representations by training the classifier to distinguish auditory from visual information. There was strong evidence (Figure 4b) that this network exhibited differential activation for the auditory and visual tasks (mean classification accuracy 79.3%, BF_10_ = 4.96*10^12^). The data for the individual MD ROIs (supplementary figure 2) show a similar pattern. To identify if there were any regions showing cross-generalisation outside of this network, we conducted an exploratory analysis using a roaming searchlight and assessed the results with cluster-level family wise error correction for multiple comparisons (voxelwise threshold: *p* < 0.005 uncorrected). No clusters survived this correction.

**Figure 4.**
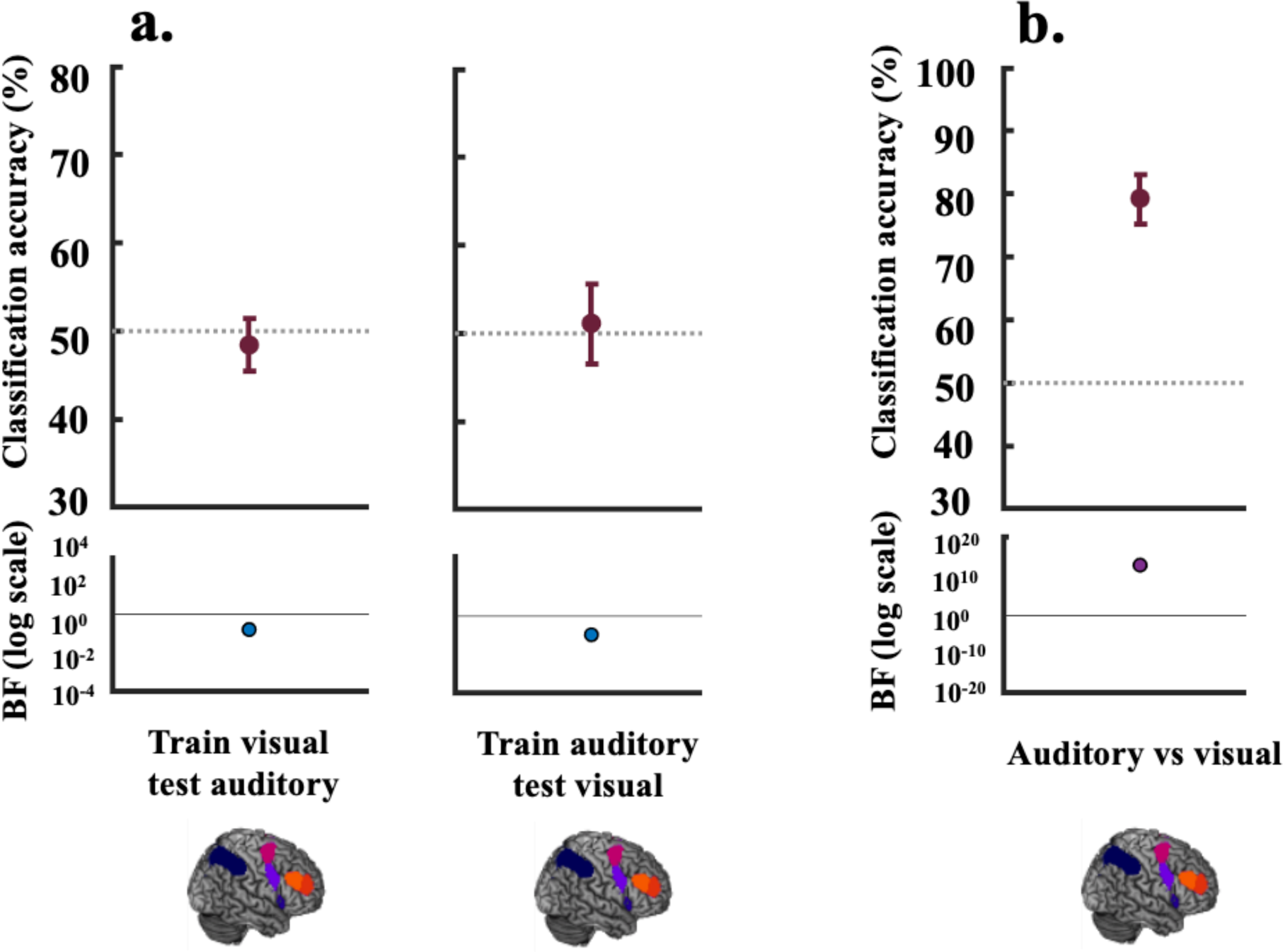
Decoding results for cross-generalisation (between modality decoding) vs modality-specific representations. For Panel a) the classifier was trained on the visual rules (visual rule 1 vs visual rule 2) and tested on the auditory rules (auditory rule 1 vs auditory rule 2) (left panel) and vice-versa (right panel). The lower parts of the plots show the associated BFs on a logarithmic scale, generated using custom code (from Teichmann et al., 2021). BF_10_ < 1/3 are marked in blue and BF_10_ > 3 are marked in purple-coloured circles. The BFs for cross-generalisation indicate evidence for the null (BF_10_ [0.16, 0.11]) that the MD network does not hold an abstract modality-independent representation of the rules with cross-generalisable underlying codes. Panel b) shows classification of the auditory (rule 1 and rule 2) vs the visual task (rule 1 and rule 2). This positive control shows evidence that this network distinguishes between information pertaining to the two modalities (BF_10_ = 4.96*10^12^). For plotting purposes, we bootstrapped 95% confidence intervals across participants using 10,000 bootstrap samples.

Finally, we performed a sanity check for our cross-generalisation classification analyses, assessing whether we could generalise motor responses in BA4 across the auditory and visual tasks. We anticipated that we should not only be able to successfully decode the given response (inner vs outer finger position) in BA4 but moreover that this information would not be encoded differentially based on sensory modality (i.e., it would cross-generalise) and this would therefore be a good test of the cross-generalisation analysis. As depicted in Figure 5 were able to decode the inner vs. outer responses in both the visual and auditory tasks, and cross-generalise between them.

**Figure 5.**
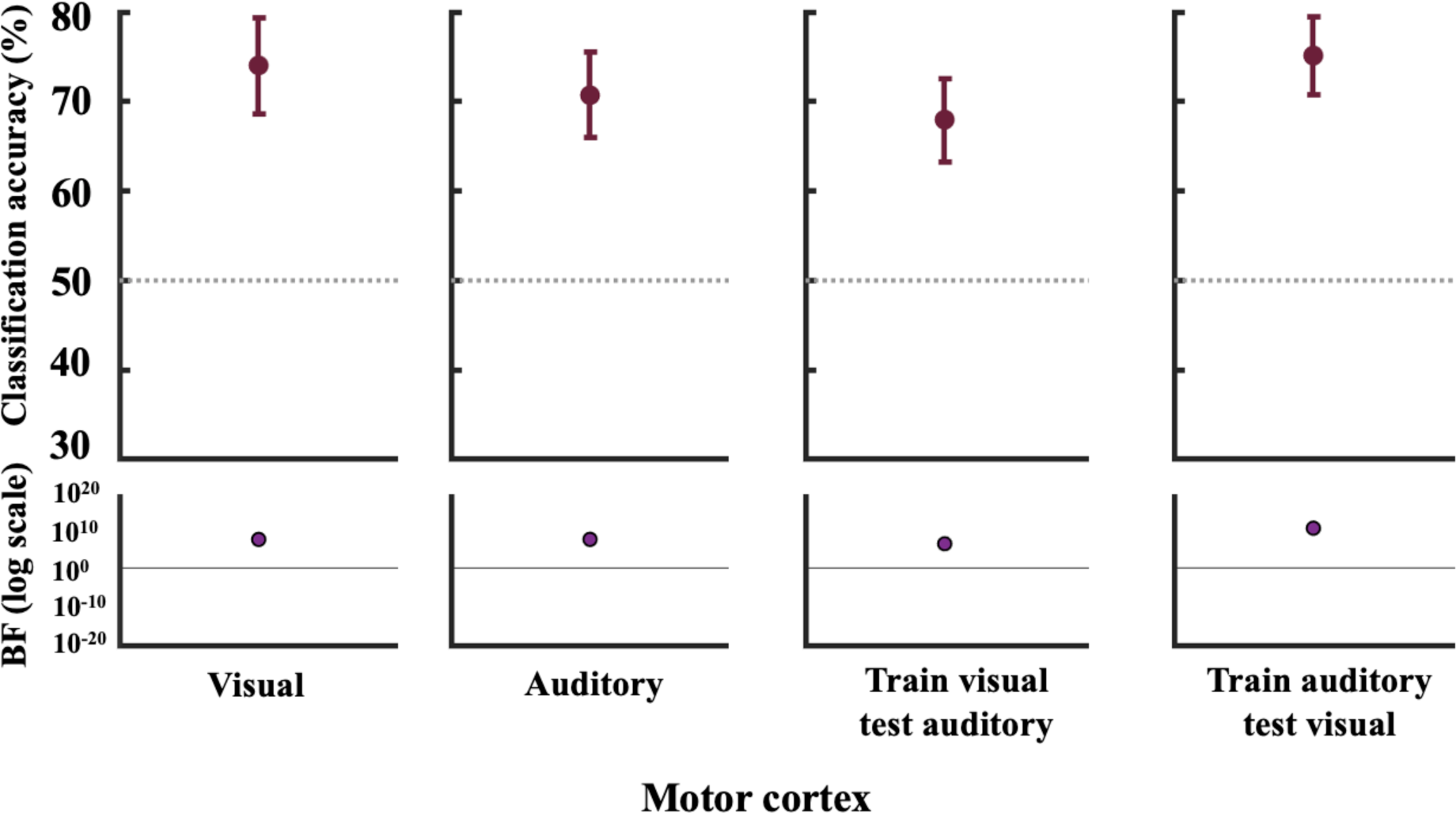
Decoding and BF results for the motor cortex. We performed a sanity check to assess whether we could decode motor responses and whether this was independent of modality. From left to right we decoded inner vs. outer finger responses 1) in the visual task 2) in the auditory task 3) training in the visual, testing in the auditory, and 4) training in the auditory, testing in the visual. For plotting purposes, we bootstrapped 95% confidence intervals across participants using 10,000 bootstrap samples. The lower parts of the plots show the associated BFs on a logarithmic scale, generated using custom code (from Teichmann et al., 2021). BF_10_ < 1/3 are marked in blue and BF_10_ > 3 are marked in purple-coloured circles.

## Discussion

In this study we wanted to understand how domain generality arises in the frontoparietal MD network, including whether information that could theoretically be abstracted away from the input modality would be coded in a modality-general abstract form. We tested 1) whether the MD network of the human brain encodes information about both auditory and visual-based rules; 2) whether the same neural resources (voxels) contribute to both sets of codes; and 3) whether the codes underlying these task-relevant rule representations are shared across modality or remain modality-specific. There was evidence that this network coded rule information for both auditory and visual tasks, corresponding with the dominant view of this network as domain general. Further, more of the most strongly responding voxels than would be expected by chance contributed to both classification schemes, suggesting that some MD resources may be re-allocated to code task information pertaining to different sensory inputs. However, there was evidence that the patterns underlying coding of auditory and visual rules did not cross-generalise. Alongside this, there was evidence from our positive control that the classifier could distinguish between the same rules applied to the two modalities. These data suggest that although this domain general network holds information pertaining to distinct sensory inputs, and the same neural resources are implicated in doing so, the codes underlying the representation of this information are independent and tagged by modality, rather than shared and abstracted to the level of the rule.

The findings from this study complement ongoing research showing that the MD network processes information from not only a wide-range of tasks, but also across modalities (Duncan & Owen, 2000; Duncan, 2010; Duncan et al., 2020; Assem et al., 2022; Schultz et al., 2022). The data also fit well with the theory that this network is key to integration of various types of information (Duncan et al., 2020). This is thought to be the case due to strong connectivity between the core regions of the network (Assem et al., 2020) and task-dependent connections to other networks (Cole & Schneider, 2007; Cole et al., 2013; Cocuzza et al., 2020). Our data add to this base by showing that not only are both visual and auditory rules encoded in these regions, but the patterns for each tend to load onto the same MD voxels.

These results showed that, even with a presumably imperfect MD definition, more voxels than would be expected by chance, contributed to the representation of both types of information (rule encoding in the auditory, and in the visual domain). By contrast, other work has shown modality biased subregions in and around the MD network (Michalka, Kong, Rosen, Shinn-Cunningham, & Somers, 2015; Braga, Hellyer, Wise, & Leech, 2017; Mayer, Ryman, Hanlon, Dodd, & Ling, 2017; Noyce, Cestero, Michalka, Shinn-Cunningham, & Somers, 2017). For example, contrasting visual and auditory attention tasks has revealed two visual-biased regions along the superior and inferior precentral sulcus (PCS), interleaved with two auditory-biased regions along the transverse gyrus intersecting the PCS and along the caudal portion of the IFS (Michalka et al., 2015; Noyce et al., 2017). Other MD regions (e.g. anterior cingulate and insula) have not shown sensory biases (Noyce et al., 2017) and using high precision fMRI methods, sensory-biased areas identified around lateral frontal cortex appear to lie adjacent to the MD borders (Assem et al., 2022). Taken together this previous work and the findings from the present study emphasise that MD neurons may differ in their relative potential for coding different types of information, and thus show sensory biases under specific circumstances, but that they are also highly adaptable (Duncan, 2001) and can therefore potentially encode many different types of information. What types of information are encoded may depend on many factors, for example the nature of the task. Our data suggests that there is a flexible allocation of neural resources between our two tasks at least, perhaps reflective of mixed selectivity (Rigotti et al., 2013). Although, it is important to note that while our voxel re-use analysis allows greater spatial specificity than looking at whole regions, we cannot draw conclusions at the neural level as independent populations may underlie these voxel responses. The current picture suggests that while there may be sub-preferences towards different types of information, MD cells are highly adaptable and can be flexibly allocated to encode different types of information, perhaps depending on the nature of the task.

While some of the voxels with the strongest signal were the same across the two classification schemes, they did not appear to be used to form similar, modality-independent, codes. Instead, there was evidence for the null: there was no cross-generalisation between the auditory rules and visual rules, indicative of independent neural representations of modality-based rule information. By contrast, motor responses were decodable in the motor cortex, and it was possible to cross-generalise motor responses between the two modality-based tasks. In domain general areas, independent representations of task rule for each modality would arise if general-processing resources randomly or systematically become associated with particular inputs and outputs, forming bespoke conjunctions. This aligns with recent recurrent models of working memory in higher cortical regions (Bouchacourt & Buschman, 2019; Manohar et al., 2019; Buschman, 2021). In these models, general-purpose “conjunction” units come to represent conjunctions of stimuli due to fixed random recurrent connections (Buschman, 2021) or flexibly through rapid Hebbian updating of synaptic weights that form bespoke conjunctions depending on the current task (Manohar et al., 2019). Accordingly, neurons in these regions have been shown to respond to a conjunction of different sensory inputs, in different contexts and at particular points in time (Mante, Sussillo, Shenoy, & Newsome, 2013; Rigotti et al., 2013; Aoi, Mante, & Pillow, 2020; Bocincova et al., 2022). These mixed-selectivity properties result in high dimensional spaces for representing cognitive variables and maintaining a unique combination of inputs, capturing the diversity of information from different brain regions (Bouchacourt & Buschman, 2019; Badre et al., 2021; Buschman, 2021). One possibility is that recurrent conjunctive coding in the MD system extends also to combinations of stimuli and responses. Moreover, if this is underpinned by a flexible association process (Manohar et al., 2019), it could provide a basis for the human ability to respond flexibly according to current task rules. This would then align with the proposal that a key role of the MD system may be to create the necessary associations between the information and actions needed for the current task (Duncan et al., 2020). Our data fit well within this framework, suggesting that the system flexibly responded here, using the same resources to form independent representations of the stimulus-response rules in the auditory and visual tasks, rather than abstract, modality-independent, codes.

The choice of task design, imaging technique, and analysis will influence how information about different modalities is processed, represented, and read out. For example, in Assem and colleagues’ (2022) work, MD hard-minus-easy visual and auditory activation maps were close to identical, suggesting that the MDs can respond to both types of information, comparable to the present data, but also suggestive that responses to control demands in auditory and visual tasks may be similar. However, not only did the task design differ (n-back working memory) to the present study, but so did the analysis, which was univariate and does not provide insight about the level of representation. Our multivariate analysis suggests that while domain general areas encode information from both modalities, the information is represented in independent neural codes. In support of this, there was also strong evidence that the MD network patterns were clearly distinguishable between auditory and visual information. This analysis informs us that the network responds differently for auditory and visual trials. However, this result could have been driven by many factors, as the two tasks were well-matched but not identical. It could, for example, be driven by differences in difficulty, even though we attempted to match difficulty with a staircase procedure. Another aspect to consider is to what extent rule decoding could have been driven by visual and auditory properties of the cues. However, this is unlikely to explain the current results as, in addition to using two cues per modality, the cue occurred a few seconds earlier than the time that we modelled, with several stimulus events occurring in between.

Our cross-generalisation analysis appears to align with the idea that MD neurons can encode multiple task features (here, modality*rule) in a nonlinear, conjunctive fashion. However, we do not know the extent of this potential nonlinearity, only that it is sufficient to prevent cross-generalisation of the fMRI data. This leaves open the possibility that some MD cells do respond to different modalities in a more linear fashion, which we were not sensitive to here. The percentage of neurons that exhibit mixed selectivity is unknown, and it may vary from one region to the next, and in any given task (Parthasarathy et al., 2017; Dang, Li, Pu, Qi, & Constantinidis, 2022). We also cannot address here whether non-generalisable coding is a general property of the MD system. It could be unique to specific features such as modality, while in other cases, task features are combined linearly. For example, following training, monkey prefrontal neurons have shown increased nonlinear mixed selectivity for task features in a spatial but not in a shape-based working memory task (Dang, Jaffe, Qi, & Constantinidis, 2021). The way that task demands change over time may also impact the level of abstraction at which representations are held. For example, again in monkey prefrontal cortex (e.g., Bernardi et al., 2020) it was shown that the level of abstraction at which various task representations are held changes before and during the course of a trial. Here, we used a low temporal resolution technique (fMRI) but future work could use neural measurements with sufficient resolution, such as magnetoencephalography, to assess how these types of representations change over time.

The MD network is widely known thought of as domain general but there are many different possible conceptualisations of what domain generality is. Here we sought to characterise domain generality in the MD system in terms of how information from multiple sensory modalities is represented. Our findings confirm that the MD network encodes information from multiple modalities, here rules pertaining to both visual and auditory tasks. As far as the resolution of fMRI permits, the data also suggested that similar neural resources were used to code for both sets of rules. However, the underling neural codes reflecting task rules in the context of each modality were not integrated to such a level of abstraction that we could cross-generalise between them. We therefore suggest that while neural resources in the MD system may be flexibly assigned to represent arbitrary conjunctions of stimuli and responses, providing a basis for the human ability to respond flexibly to what we see and hear, this ability does not rely on forming abstract representations of rules that generalise over input modalities.

## Author contributions

Conceptualisation: A.N.R., J.D., and A.W.; investigation: L.T., D.M.; formal analysis: J.B.J., D.M., L.T., and A.W.; writing—original draft: J.B.J.; writing—review and editing: J.B.J., L.T., D.M., J.D., A.N.R. and A.W.; funding acquisition: A.N.R. and A.W.

## Data availability

The ethical approval for this study does not allow us to share raw data openly. Source data for main report Figs. 2-5 and supplementary Figs. 1-3 is available on Open Science Framework (OSF) (osf.io/hcpku). Template regions of interest and the code used to analyse the current study are publicly available on OSF (osf.io/hcpku).

## Acknowledgements

This project was supported by Australian Research Council (ARC) Discovery Project Grants to A.W., A.R. and J.D (DP170101840; DP120102835). J.J. and A.W. are currently supported by MRC (U.K) intramural funding SUAG/093/G116768. For the purpose of open access, the author has applied a Creative Commons Attribution (CC BY) licence to any Author Accepted Manuscript version arising from this submission.

## References

Aoi, M. C., Mante, V., & Pillow, J. W. (2020). Prefrontal cortex exhibits multidimensional dynamic encoding during decision-making. Nature neuroscience, 23(11), 1410–1420.

Assem, M., Glasser, M. F., Van Essen, D. C., & Duncan, J. (2020). A domain-general cognitive core defined in multimodally parcellated human cortex. Cerebral Cortex, 30(8), 4361–4380.

Assem, M., Shashidhara, S., Glasser, M. F., & Duncan, J. (2022). Precise topology of adjacent domain-general and sensory-biased regions in the human brain. Cerebral Cortex, 32(12), 2521–2537.

Azuma, M., & Suzuki, H. (1984). Properties and distribution of auditory neurons in the dorsolateral prefrontal cortex of the alert monkey. Brain research, 298(2), 343–346.

Badre, D., Bhandari, A., Keglovits, H., & Kikumoto, A. (2021). The dimensionality of neural representations for control. Current Opinion in Behavioral Sciences, 38, 20–28.

Bernardi, S., Benna, M. K., Rigotti, M., Munuera, J., Fusi, S., & Salzman, C. D. (2020). The geometry of abstraction in the hippocampus and prefrontal cortex. Cell, 183(4), 954–967. e921.

Bocincova, A., Buschman, T. J., Stokes, M. G., & Manohar, S. G. (2022). Neural signature of flexible coding in prefrontal cortex. Proceedings of the National Academy of Sciences, 119(40), e2200400119.

Bouchacourt, F., & Buschman, T. J. (2019). A flexible model of working memory. Neuron, 103(1), 147–160. e148.

Braga, R. M., Hellyer, P. J., Wise, R. J., & Leech, R. (2017). Auditory and visual connectivity gradients in frontoparietal cortex. Human Brain Mapping, 38(1), 255–270.

Buschman, T. J. (2021). Balancing flexibility and interference in working memory. Annual review of vision science, 7, 367–388.

Calhoun, V. D., Stevens, M. C., Pearlson, G. D., & Kiehl, K. A. (2004). fMRI analysis with the general linear model: removal of latency-induced amplitude bias by incorporation of hemodynamic derivative terms. Neuroimage, 22(1), 252–257.

Cocuzza, C. V., Ito, T., Schultz, D., Bassett, D. S., & Cole, M. W. (2020). Flexible coordinator and switcher hubs for adaptive task control. Journal of Neuroscience, 40(36), 6949–6968.

Cole, M. W., Reynolds, J. R., Power, J. D., Repovs, G., Anticevic, A., & Braver, T. S. (2013). Multi-task connectivity reveals flexible hubs for adaptive task control. Nature neuroscience, 16(9), 1348–1355.

Cole, M. W., & Schneider, W. (2007). The cognitive control network: integrated cortical regions with dissociable functions. Neuroimage, 37(1), 343–360.

Dang, W., Jaffe, R. J., Qi, X.-L., & Constantinidis, C. (2021). Emergence of nonlinear mixed selectivity in prefrontal cortex after training. Journal of Neuroscience, 41(35), 7420–7434.

Dang, W., Li, S., Pu, S., Qi, X.-L., & Constantinidis, C. (2022). More prominent nonlinear mixed selectivity in the dorsolateral prefrontal than posterior parietal cortex. Eneuro, 9(2).

Dienes, Z. (2011). Bayesian versus orthodox statistics: Which side are you on? Perspectives on Psychological Science, 6(3), 274–290.

Duncan, J. (2001). An adaptive coding model of neural function in prefrontal cortex. Nature Reviews Neuroscience, 2(11).

Duncan, J. (2010). The multiple-demand (MD) system of the primate brain: mental programs for intelligent behaviour. Trends in cognitive sciences, 14(4), 172–179.

Duncan, J., Assem, M., & Shashidhara, S. (2020). Integrated intelligence from distributed brain activity. Trends in Cognitive Sciences.

Duncan, J., & Owen, A. M. (2000). Common regions of the human frontal lobe recruited by diverse cognitive demands. Trends in neurosciences, 23(10), 475–483.

Fedorenko, E., Duncan, J., & Kanwisher, N. (2013). Broad domain generality in focal regions of frontal and parietal cortex. Proceedings of the National Academy of Sciences, 110(41), 16616–16621.

Freedman, D. J., & Assad, J. A. (2006). Experience-dependent representation of visual categories in parietal cortex. Nature, 443(7107), 85–88.

Freedman, D. J., Riesenhuber, M., Poggio, T., & Miller, E. K. (2001). Categorical representation of visual stimuli in the primate prefrontal cortex. Science, 291(5502), 312–316.

Haufe, S., Meinecke, F., Gorgen, K., Dahne, S., Haynes, J. D., Blankertz, B., & Biessmann, F. (2014). On the interpretation of weight vectors of linear models in multivariate neuroimaging. Neuroimage, 87, 96–110. doi:10.1016/j.neuroimage.2013.10.067

Hebart, M. N., Görgen, K., & Haynes, J.-D. (2015). The Decoding Toolbox (TDT): a versatile software package for multivariate analyses of functional imaging data. Frontiers in neuroinformatics, 8, 88.

Hemerik, J., & Goeman, J. (2018). Exact testing with random permutations. Test, 27(4), 811–825.

Jackson, J., Rich, A. N., Williams, M. A., & Woolgar, A. (2017). Feature-selective attention in frontoparietal cortex: multivoxel codes adjust to prioritize task-relevant information. Journal of cognitive neuroscience, 29(2), 310–321.

Jackson, J. B., Feredoes, E., Rich, A. N., Lindner, M., & Woolgar, A. (2021). Concurrent neuroimaging and neurostimulation reveals a causal role for dlPFC in coding of task-relevant information. Communications biology, 4(1), 588.

Jackson, J. B., & Woolgar, A. (2018). Adaptive coding in the human brain: Distinct object features are encoded by overlapping voxels in frontoparietal cortex. cortex, 108, 25–34.

Jeffreys, H. (1998). The theory of probability: OUP Oxford.

Jenkinson, M., Beckmann, C. F., Behrens, T. E., Woolrich, M. W., & Smith, S. M. (2012). Fsl. Neuroimage, 62(2), 782–790.

Ji, J. L., Spronk, M., Kulkarni, K., Repovš, G., Anticevic, A., & Cole, M. W. (2019). Mapping the human brain’s cortical-subcortical functional network organization. Neuroimage, 185, 35–57.

Jiang, Y., & Kanwisher, N. (2003). Common neural substrates for response selection across modalities and mapping paradigms. Journal of cognitive neuroscience, 15(8), 1080–1094.

Kass, R. E., & Raftery, A. E. (1995). Bayes factors. Journal of the american statistical association, 90(430), 773–795.

Kriegeskorte, N., Goebel, R., & Bandettini, P. (2006). Information-based functional brain mapping. Proceedings of the National academy of Sciences of the United States of America, 103(10), 3863–3868.

Manohar, S. G., Zokaei, N., Fallon, S. J., Vogels, T. P., & Husain, M. (2019). Neural mechanisms of attending to items in working memory. Neuroscience & Biobehavioral Reviews, 101, 1–12.

Mante, V., Sussillo, D., Shenoy, K. V., & Newsome, W. T. (2013). Context-dependent computation by recurrent dynamics in prefrontal cortex. nature, 503(7474), 78–84.

Mayer, A. R., Ryman, S. G., Hanlon, F. M., Dodd, A. B., & Ling, J. M. (2017). Look hear! The prefrontal cortex is stratified by modality of sensory input during multisensory cognitive control. Cerebral Cortex, 27(5), 2831–2840.

Michalka, S. W., Kong, L., Rosen, M. L., Shinn-Cunningham, B. G., & Somers, D. C. (2015). Short-term memory for space and time flexibly recruit complementary sensory-biased frontal lobe attention networks. Neuron, 87(4), 882–892.

Morey, R. D., Romeijn, J.-W., & Rouder, J. N. (2016). The philosophy of Bayes factors and the quantification of statistical evidence. Journal of Mathematical Psychology, 72, 6–18.

Noyce, A. L., Cestero, N., Michalka, S. W., Shinn-Cunningham, B. G., & Somers, D. C. (2017). Sensory-biased and multiple-demand processing in human lateral frontal cortex. Journal of Neuroscience, 37(36), 8755–8766.

Parthasarathy, A., Herikstad, R., Bong, J. H., Medina, F. S., Libedinsky, C., & Yen, S.-C. (2017). Mixed selectivity morphs population codes in prefrontal cortex. Nature neuroscience, 20(12), 1770–1779.

Peelle, J. E. (2014). Methodological challenges and solutions in auditory functional magnetic resonance imaging. Frontiers in neuroscience, 8, 253.

Peelle, J. E., Eason, R. J., Schmitter, S., Schwarzbauer, C., & Davis, M. H. (2010). Evaluating an acoustically quiet EPI sequence for use in fMRI studies of speech and auditory processing. Neuroimage, 52(4), 1410–1419.

Perrachione, T. K., & Ghosh, S. S. (2013). Optimized design and analysis of sparse-sampling fMRI experiments. Frontiers in neuroscience, 7, 55.

Petit, S., Badcock, N. A., Grootswagers, T., Rich, A. N., Brock, J., Nickels, L., Moerel, D., Dermody, N., Yau, S., & Schmidt, E. (2020). Toward an individualized neural assessment of receptive language in children. Journal of Speech, Language, and Hearing Research, 63(7), 2361–2385.

Phipson, B., & Smyth, G. K. (2010). Permutation P-values should never be zero: calculating exact P-values when permutations are randomly drawn. Statistical applications in genetics and molecular biology, 9(1).

Pischedda, D., Görgen, K., Haynes, J.-D., & Reverberi, C. (2017). Neural representations of hierarchical rule sets: the human control system represents rules irrespective of the hierarchical level to which they belong. Journal of Neuroscience, 37(50), 12281–12296.

Power, J. D., Cohen, A. L., Nelson, S. M., Wig, G. S., Barnes, K. A., Church, J. A., Vogel, A. C., Laumann, T. O., Miezin, F. M., & Schlaggar, B. L. (2011). Functional network organization of the human brain. Neuron, 72(4), 665–678.

Rao, S. C., Rainer, G., & Miller, E. K. (1997). Integration of what and where in the primate prefrontal cortex. Science, 276(5313), 821–824.

Rigotti, M., Barak, O., Warden, M. R., Wang, X.-J., Daw, N. D., Miller, E. K., & Fusi, S. (2013). The importance of mixed selectivity in complex cognitive tasks. Nature, 497(7451), 585–590.

Robinson, A. K., Rich, A. N., & Woolgar, A. (2022). Linking the brain with behavior: the neural dynamics of success and failure in goal-directed behavior. Journal of cognitive neuroscience, 34(4), 639–654.

Romanski, L. M. (2007). Representation and integration of auditory and visual stimuli in the primate ventral lateral prefrontal cortex. Cerebral Cortex, 17(suppl_1), i61–i69.

Rorden, C. (2007). Mricron [computer software].

Rouder, J. N., Speckman, P. L., Sun, D., Morey, R. D., & Iverson, G. (2009). Bayesian t tests for accepting and rejecting the null hypothesis. Psychonomic bulletin & review, 16(2), 225–237.

Schultz, D. H., Ito, T., & Cole, M. W. (2022). Global connectivity fingerprints predict the domain generality of multiple-demand regions. Cerebral Cortex, 32(20), 4464–4479.

Shashidhara, S., Spronkers, F. S., & Erez, Y. (2020). Individual-subject functional localization increases Univariate activation but not multivariate pattern discriminability in the “multiple-demand” frontoparietal network. Journal of cognitive neuroscience, 32(7), 1348–1368.

Stelzer, J., Chen, Y., & Turner, R. (2013). Statistical inference and multiple testing correction in classification-based multi-voxel pattern analysis (MVPA): random permutations and cluster size control. Neuroimage, 65, 69–82.

Team, J. (2024). JASP (Version 0.18.3)[Computer software].

Teichmann, L., Moerel, D., Baker, C., & Grootswagers, T. (2021). An empirically-driven guide on using Bayes Factors for M/EEG decoding. bioRxiv.

Todd, M. T., Nystrom, L. E., & Cohen, J. D. (2013). Confounds in multivariate pattern analysis: Theory and rule representation case study. NeuroImage, 77, 157–165. doi:10.1016/j.neuroimage.2013.03.039

Vaidya, A. R., Jones, H. M., Castillo, J., & Badre, D. (2021). Neural representation of abstract task structure during generalization. ELife, 10, e63226.

Wetzels, R., Matzke, D., Lee, M. D., Rouder, J. N., Iverson, G. J., & Wagenmakers, E.-J. (2011). Statistical evidence in experimental psychology: An empirical comparison using 855 t tests. Perspectives on Psychological Science, 6(3), 291–298.

Woolgar, A., Afshar, S., Williams, M. A., & Rich, A. N. (2015). Flexible coding of task rules in frontoparietal cortex: an adaptive system for flexible cognitive control. Journal of Cognitive Neuroscience, 27(10), 1895–1911.

Woolgar, A., Dermody, N., Afshar, S., Williams, M. A., & Rich, A. N. (2019). Meaningful patterns of information in the brain revealed through analysis of errors. bioRxiv, 673681.

Woolgar, A., Hampshire, A., Thompson, R., & Duncan, J. (2011). Adaptive coding of task-relevant information in human frontoparietal cortex. Journal of Neuroscience, 31(41), 14592–14599.

Woolgar, A., Jackson, J., & Duncan, J. (2016). Coding of visual, auditory, rule, and response information in the brain: 10 years of multivoxel pattern analysis. Journal of cognitive neuroscience.

Woolgar, A., Williams, M. A., & Rich, A. N. (2015). Attention enhances multi-voxel representation of novel objects in frontal, parietal and visual cortices. Neuroimage, 109, 429–437.

Woolgar, A., & Zopf, R. (2017). Multisensory coding in the multiple-demand regions: vibrotactile task information is coded in frontoparietal cortex. Journal of Neurophysiology, 118(2), 703–716.

