## Supplementary materials for "Domain general frontoparietal regions show modality-dependent coding of auditory and visual rules"

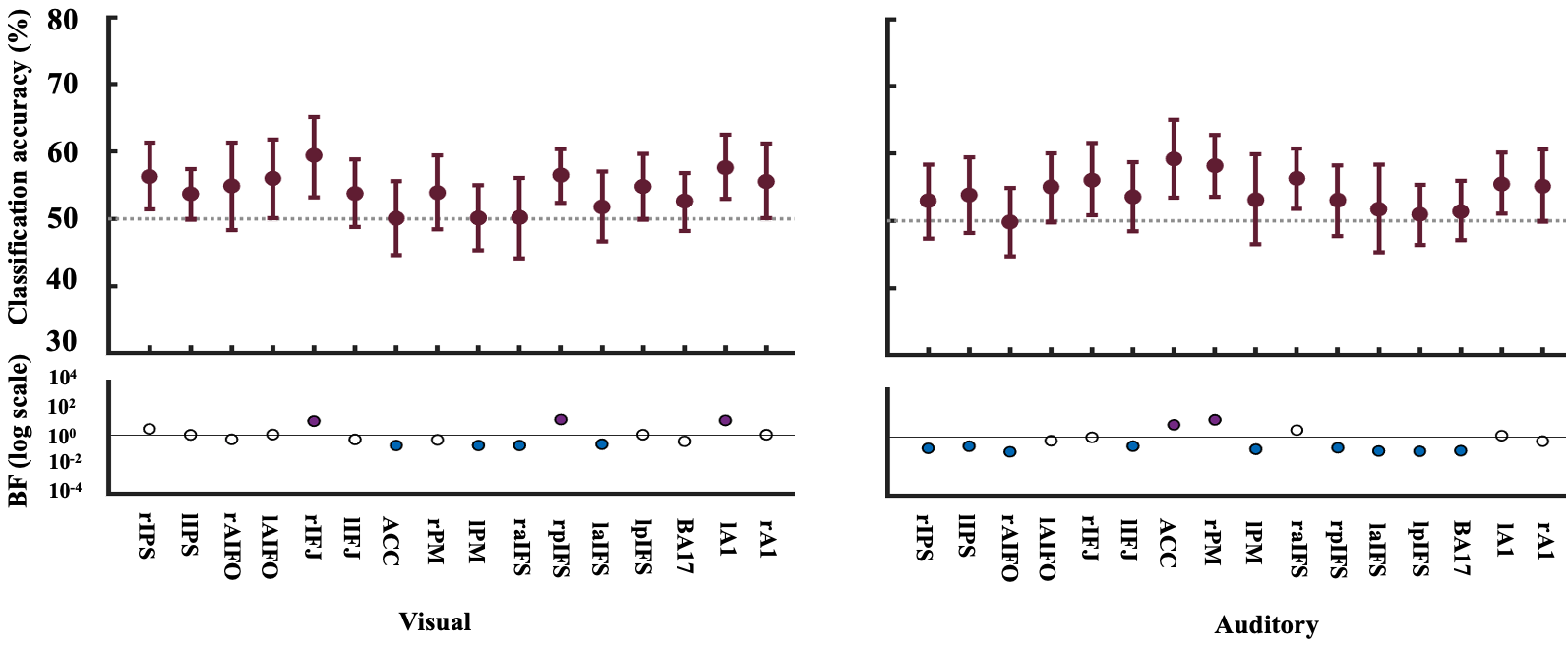

***Supplementary figure 1.*** *Decoding results for individual MD ROIs, visual and auditory cortex. On the left panel is the classification (%) and BF results for the visual rule and on the right panel is the classification (%) and BF results for the auditory rule. For plotting purposes, we bootstrapped 95% confidence intervals across participants using 10,000 bootstrap samples. The lower parts of the plots show the associated BFs on a logarithmic scale, generated using custom code (from Teichmann et al., 2021). BF*_10_ *< 1/3 are marked in blue and BF*_10_ *> 3 are marked in purple-coloured circles. Similar to the combined MD ROI analysis (main manuscript), individual MDs appear to hold information about both the auditory and visual rules. These plots indicate some regional differences where there is evidence that the area holds information about the visual rule but not the auditory rule, and vice versa. Surprisingly, there is evidence that the left auditory cortex encodes information about the rules in the visual domain, otherwise the decoding results for the specialised cortices are generally inconclusive.*

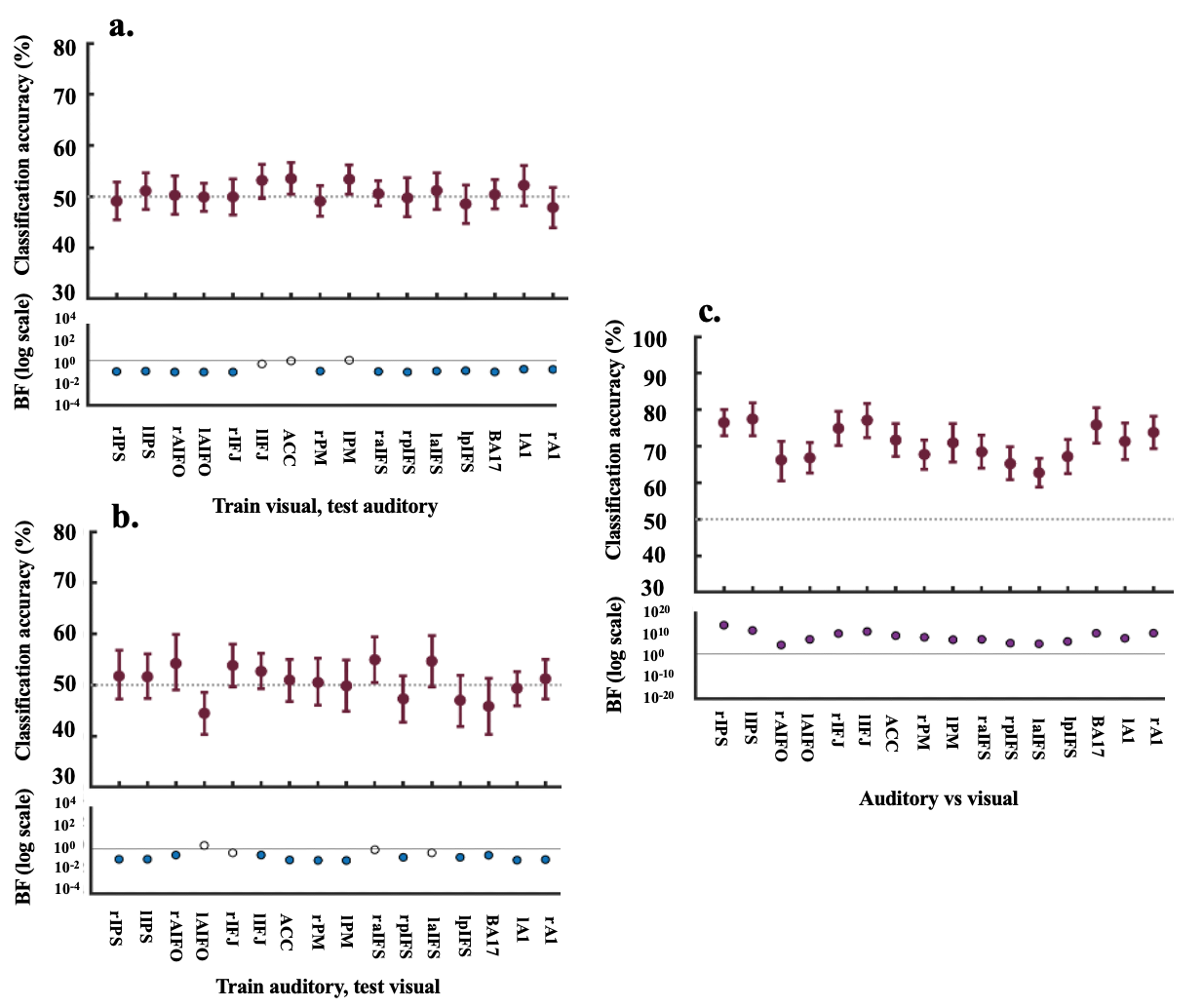

***Supplementary figure 2.*** *Decoding and BF results for individual MD ROIs, visual and auditory cortex for cross-generalisation vs modality-specific representation. For Panel a) the classifier was trained on the visual rules (visual rule 1 vs visual rule 2) and tested on the auditory rules (auditory rule 1 vs auditory rule 2) (upper) and vice-versa (Panel b). Panel c) shows classification of the auditory rules (rule 1 and rule 2) vs the visual rules (rule 1 and rule 2). For plotting purposes, we bootstrapped 95% confidence intervals across participants using 10,000 bootstrap samples. The lower parts of the plots show the associated BFs on a logarithmic scale, generated using custom code (from Teichmann et al., 2021). BF*_10_ *< 1/3 are marked in blue and BF*_10_ *> 3 are marked in purple-coloured circles. Similar to the combined MD ROI analysis (main manuscript), there is evidence for the null that there is no cross-generalisation of information between the two modalities. Instead, there is evidence in each MD ROI that we can distinguish between auditory and visual information.*
